# Cas9 fusions for precision *in vivo* editing

**DOI:** 10.1101/2020.07.15.199620

**Authors:** Ryan R. Richardson, Marilyn Steyert, Saovleak N. Khim, Andrea J. Romanowski, Jeffrey Inen, Bekir Altas, Alexandros Poulopoulos

## Abstract

Cas9 targets genomic loci with high specificity. When used for knockin, however, Cas9 often leads to unintended on-target knockout rather than intended edits. This imprecision is a barrier for direct *in vivo* editing where clonal selection is not feasible. Here we demonstrate a high-throughput workflow to ratiometrically assess on-target efficiency and precision of editing outcomes. Using this workflow, we screened combinations of donor DNA and Cas9 variants, as well as fusions to DNA repair proteins. This yielded novel high-performance double-strand break repair editing agents and combinatorial optimizations with orders-of-magnitude increases in knockin precision. Cas9-RC, a novel Cas9 fusion to eRad18 and CtIP, increased knockin performance over 3-fold *in vitro* and *in vivo* in the developing mouse brain. Continued comparative assessment of existing and novel editing agents with this ratiometric framework of efficiency and precision will further the development of direct *in vivo* knockin and future genetic therapies.

## Introduction

CRISPR technologies have enabled the development of agents to edit the genome in naive host cells and whole organisms. The development of high-performance *in vivo* editing agents offers the potential for cost- and time-effective knockin experiments in wildtype organisms without the need for transgenesis and animal lines. The broad efficacy of Cas9 across biological systems holds further promise for knockin research in non-model organisms^1^ and future genome therapeutics in humans.^2^

A key challenge for genome editing *in vivo* is outcome precision.^3^ Existing tools offer adequate precision for *ex vivo* editing, where correctly edited cells can be selected, expanded, and validated into single-clone populations. To achieve reliable editing directly *in vivo*, where selection is typically impractical, editing agents must perform with high efficiency as well as high precision to minimize unintended on-target outcomes. Developing editing agents with high performance in both efficiency and precision remains a challenge for the field.

The Cas9 toolkit offers a variety of editing modalities, including double-strand break (DSB) repair-based editing, base editing, and reverse transcriptase-based (Prime) editing.^4^ DSB repair-based approaches are maximally versatile for knockin, allowing insertions from DNA donors that are large enough for fusion-protein knockins. However, such approaches are also the most imprecise often leading to unintended disruptions in gene function.^5^ Current DSB repair-based knockin methods produce the desired editing outcome the minority of the time, an order-of-magnitude less than unintended on-target insertions or deletions (indels).^3^

Here we present a high-throughput workflow to quantify editing outcomes for creating and identifying high-performance knockin tools and their optimal combinations. We established editing *efficiency* and *precision* as generalizable assessment metrics for comparisons of knockin performance across existing and novel agents. Using this platform, we aimed to enhance DSB repair-based editing performance by combinatorial screens of Cas9 variants, DNA donors, and novel compound fusions to DNA repair proteins. This workflow yielded Cas9-RC, a novel high-performance DSB repair-based editing agent with increased editing efficiency and precision. We tested Cas9-RC for its editing performance *in vivo* in the embryonic mouse brain, where it enhanced fluorescent protein knockin by 370%. These improvements showcase the utility of this workflow for continued development and assessment of new precision *in vivo* editing tools.

## Methods

### Plasmid design and construction

Mammalian expression plasmids and knockin donor template plasmids were constructed with a combination of standard cloning techniques. For gRNA expression constructs, oligos (Integrated DNA Technologies) were annealed and cloned into a custom hU6 backbone using Golden Gate Assembly (GGA).^6^ Cas9 expression constructs were assembled by a modified mMoClo system.^7^ Briefly, individual parts were cloned, BsaI adapters added, and internal sites removed via PCR using KAPA HiFi HotStart DNA Polymerase with 2X Master Mix (Roche), or synthesized (Integrated DNA Technologies). Parts were subsequently assembled into expression constructs using NEB Golden Gate Assembly Kit (BsaI-HF v2) according to manufacturer’s recommendations. Homology arms for donor templates were PCR amplified from CD1 mouse genomic DNA with adapters for GGA. gRNA binding site (GRBS) parts were generated by oligo annealing. GRBS and homology arms were assembled with knockin sequences via GGA. See Supplementary Table S1 for further detail.

### Cell culture and transfection

HEK:BFP cells were the kind gift of Chris Richardson.^8^ Cells were maintained at 37°C and 5% CO2 in DMEM media plus GlutaMax (ThermoFisher Scientific) supplemented with 10% (v/v) fetal bovine serum (FBS). Typically, 20,000-22,500 cells/cm2 were seeded onto 24-well plates the day before transfection. Cells were transiently transfected at 70-80% confluence using Polyethylenimine, Linear, MW 25000 (‘PEI’, Polysciences) resuspended to 1mg/mL in H2O at a 3:1 (v/w) PEI:DNA ratio with 250 ng DNA per plasmid (750 ng total DNA) diluted in Opti-MEM (ThermoFisher Scientific) and added dropwise to cells.

### Flow cytometry

Cells were trypsinized, pelleted, and resuspended in Dulbecco’s PBS containing 0.1% FBS. At least 20,000 live cells (typically 80,000+) were analyzed using an LSRII cell analyzer with HTS (BD Biosciences). BFP and mTagBFP were measured with a 407 nm laser and a 450/50 emission filter. GFP and mNeonGreen were measured with a 488 nm laser, a 505 LP mirror, and a 530/30 emission filter. mCherry was measured with a 561 nm laser, a 600 LP mirror, and a 615/25 emission filter. Data were analyzed with FlowJo v10.6.2 (Flowjo LLC). Live cells were gated by size and granularity using FSC-A vs SSC-A. Singlets were gated using SSC-A vs SSC-H (see Supplementary Fig. S1). At least 3 biological replicates were run with internal technical duplicates or triplicates.

### Sequence analysis of knockin products

Transiently transfected HEK:BFP cells were sorted using a FACSAria II sorter (BD Biosciences). Genomic DNA was extracted from sorted cells using the Genomic DNA Clean & Concentrator kit (Zymo Research). PCR fragments were amplified using KAPA HiFi HotStart DNA Polymerase with 2X Master Mix (Roche), gel extracted with Zymoclean Gel DNA Recovery kit (Zymo Research), and submitted for Sanger sequencing (Genewiz). See Supplementary Table S1 for primer sequences. Alignment of sequencing results was performed using Benchling (https://benchling.com). Analysis of editing outcomes by decomposition of Sanger sequencing data was performed using the ICE Analysis tool v2 (Synthego) as previously described.^9^

### Animals

All animal experimental protocols were approved by the University of Maryland Baltimore Institutional Animal Care and Use Committee and complied with all relevant ethical regulations regarding animal research. Experiments were performed on outbred strain CD1 mouse pups (Charles River Laboratories). Analyses are thought to include animals of both sexes at approximately equal proportions, as no sex determination was attempted. No statistical methods were used to predetermine sample size.

### *In utero* electroporation

Electroporations of plasmid DNA were performed in utero on embryonic day 14.5 (E14.5) to target cortical layer II/III, as previously described.^10^,^11^ Briefly, DNA solutions were prepared to 4 μg/μL total DNA, with 1 μg/μL of each of the relevant plasmids (donor, guide, Cas9, and fluorescent protein). Dames were deeply anesthetized with isoflurane under a vaporizer with thermal support (Patterson Scientific Link7 & Heat Therapy Pump HTP-1500). The abdominal area was prepared for surgery with hair removal, surgical scrub, and 70% ethanol and 10% Betadine solution. A midline incision was made to expose the uterine horns. Using pulled (Narishige PC-100) and beveled (Narishige EG-45) glass micropipettes connected to a pneumatic aspirator, DNA solution was injected into one lateral brain ventricle, followed by application of 4 x 50 ms square pulses of 35 V (NEPA21 electro-kinetic platinum tweezertrodes connected to a BTX ECM-830 electroporator) targeting nascent sensorimotor areas of the cortical plate. Typically, 4-6 pups were electroporated per dame. Uterine horns were placed back inside the abdominal cavity, and monofilament nylon sutures (AngioTech) were used to close muscle and skin incisions. After term birth, electroporated mouse pups were non-invasively screened for unilateral cortical fluorescence using a fluorescence stereoscope (Leica MZ10f with X-Cite FIRE LED light source) and returned to their dame until postnatal day 7 (P7). When possible, to minimize inter-dame variation, control and experimental electroporations were performed in littermate pups from the same dame.

### Histology and immunolabeling

Tissue was prepared by intracardial perfusion with PBS and 4% paraformaldehyde. Brains were cut to 80 μm coronal sections on a vibrating microtome (Leica VT1000). Sections were immunolabeled in blocking solution consisting of 5% bovine serum albumin and 0.2% Triton X-100 in PBS for 30 minutes, then incubated overnight at 4°C with primary antibodies diluted in blocking solution. Sections were washed in PBS, incubated for 3h at room temperature with secondary antibodies diluted 1:1000 in blocking solution. Following PBS washes, sections were mounted on slides with Fluoromount-G Mounting Medium with DAPI (ThermoFisher Scientific). For antibodies used, see *Antibodies* tab in Supplementary Table S1.

### Microscopy and image analysis

Fluorescence images were acquired using a Nikon Ti2-E inverted microscope fitted with an automated registered linear motor stage (HLD117, Pior Scientific), a Spectra-X 7 channel LED light engine (Lumencor), and standard filter sets for DAPI, FITC, TRITC, and Cy5. Images were analyzed with NIS-Elements (Nikon) using an automated script to identify and count electroporated cells in brain sections. mCherry-positive (mCherry^+^) knockin neurons were counted manually using ImageJ (NIH) and independently by three blinded investigators. Five 80 μm sections, centered at the middle of the anteroposterior axis of the electroporation field and taken every other section, were analyzed per brain, with counts aggregated across sections from the same brain.

### Statistical analysis

All statistical values are presented as mean ± SEM. For experiments containing more than two conditionals or groups, statistical significance was calculated using a one-way ANOVA with Tukey’s multiple comparison test, with a single pooled variance. For experiments containing two conditionals or groups, statistical significance was calculated using a two-tailed Student’s T test. Effect size was calculated using Cohen’s *d* and expressed in pooled standard deviations.^12^ Differences between conditions were judged to be significant at P < 0.05 (*), P < 0.01 (**), and P < 0.001 (***).

### Citation diversity statement

Toward awareness and mitigation of citation biases, we used cleanBib (https://github.com/dalejn/cleanBib) to assess the predicted gender and predicted racial/ethnic category of the first and last author of our cited references. The gender breakdown of our references is 11.05% woman(first)/woman(last), 9.28% man/woman, 22.18% woman/man, and 57.49% man/man. The racial/ethnic breakdown of our references is 24.39% author of color (first)/author of color(last), 13.8% white author/author of color, 31.36% author of color/white author, and 30.45% white author/white author. These analyses exclude self-citations and are subject to the caveats and limitations outlined in the cleanBib documentation.

## Results

### Establishing genomic BFP-to-GFP platform for high-throughput assessment of editing efficiency and precision

To quantify and compare Cas9 and donor variant combinations on knockin performance, we used a BFP-to-GFP conversion assay in an engineered HEK293 cell line (HEK:BFP) genomically expressing BFP.^8^ Here, donor templates target the BFP sequence and convert it to GFP via introduction of the point mutation H26Y. Fluorescence was used as a surrogate for editing outcomes following transfection of these cells with Cas9, gRNA targeting the *BFP* locus, and H26Y donor DNA. Precise H26Y knockin would cause fluorescence to shift from BFP to GFP, while on-target indels by error-prone repair would result in unintended knockout and loss of fluorescence (Fig. 1A). By quantifying the proportions of BFP^+^, GFP^+^, and dark (BFP^−^/GFP^−^) cells, we can ratiometrically assess knockin efficiency (% of GFP^+^ to BFP^+^ cells) and knockin precision (ratio of % GFP^+^ to % dark cells) across editing agents.

**Fig. 1.**
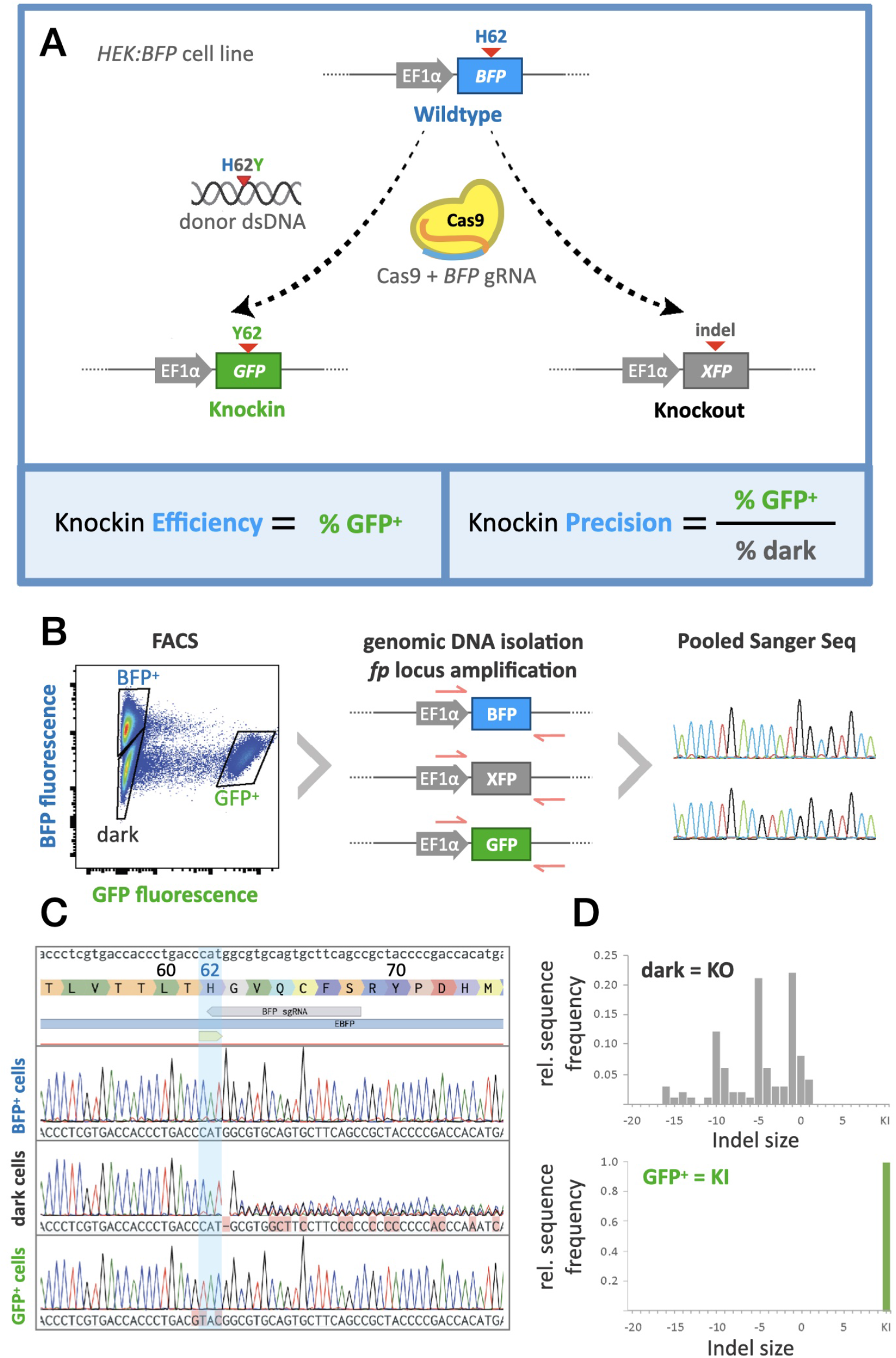
BFP-to-GFP editing as a platform to quantify knockin efficiency and precision. **(A)** Schematic of BFP-to-GFP conversion assay screen. Genome editing agents are tested by transfecting *HEK:BFP* cells, a HEK293 knockin cell line that genomically expresses BFP driven by an EF1α promoter. Successful editing knocks in the H62Y mutation into the *BFP* locus, thus producing GFP. Precisely edited cells will change from blue to green, while imprecise edits on the *BFP* locus will largely result in indel knockouts and disrupt fluorescence, turning those cells from blue to dark. Quantification of knockin efficiency is calculated by the proportion of GFP+ cells over total, and knockin precision is calculated by the proportion of GFP+ cells over dark cells. **(B)** Schematic outlining sorting and sequencing of cells treated with agents to edit the *BFP* locus. Fluorescence plot shows FACS isolation of distinct cell populations following treatment for H62Y editing. GFP fluorescence is shown on the x-axis (log), and BFP fluorescence on the y-axis (log). Blue, green, and dark collection gates are indicated. Genotyping of the three collected populations was performed by PCR of the *BFP* locus from genomic DNA, followed by Sanger sequencing of amplified fragment pools. **(C)** Alignment of Sanger sequencing reads from BFP^+^, dark, and GFP^+^ cell populations to the reference *BFP* locus sequence (top), showing wild-type, knockout (indel), and knockin genotypes, respectively. **(D)**Editing outcomes (indels, knockin frequency) were quantified by decomposition of Sanger sequencing reads using the ICE algorithm and plotted on relative histograms binned by indel size. GFP^+^ cells represent true knockins, and over 90% of dark cells contain indels predicted to cause knockout.

To validate our workflow and confirm that fluorescence readouts correspond to predicted genotypes in HEK:BFP cells, we used FACS to sort and collect the three phenotypic cell populations (BFP^+^, GFP^+^, and dark) that emerge following treatment with BFP/H26Y editing agents. Genomic DNA was extracted from sorted cell populations and used as template in PCR with primers flanking the region targeted by the BFP gRNA within the *BFP* locus, thereby amplifying all alleles regardless of on-target editing outcome. Pooled PCR amplicons from each sorted cell population were analyzed by Sanger sequencing (Fig. 1B).

As expected, the genotype of the BFP^+^ population matched that of the WT *BFP* sequence. The dark population exhibited a complex mixture of sequence results in the vicinity of the BFP gRNA cleavage site, representing unintended on-target edits due to imprecise repair. Sequencing decomposition using the ICE algorithm^13^ on amplicons from the dark sorted population revealed a predominance of deleterious indels (79% frameshift vs. 11% in-frame), in line with a loss of fluorescence due to knockout. Finally, the GFP^+^ population exhibited a single genotype containing the desired H26Y point mutation knockin from the donor template (Fig. 1C-D). The sequencing results matched the fluorescence surrogates, thus validating this platform for high-throughput ratiometric screening of knockin agents.

### Combinatorial screening of CRISPR knockin elements for improved performance

We sought to identify elements of DSB repair knockin agents that in combination offer improved efficiency and precision of editing. We investigated a matrix of combinations of three factors, which have individually been shown to enhance knockin performance: i) wildtype *S. pyogenes* Cas9 (‘WT’ Cas9) versus a High-Fidelity (‘HF’) Cas9 sequence variant (Cas9-HF);^14,15^ ii) Cas9 variants alone versus fusions with the “HDR Enhancing” (HE) N-terminal fragment (1-296) of the DNA repair protein CtIP;^16,17^ iii) circular DNA donor (‘HR’) predicted to favor homologous recombination vs. *in situ* linearized DNA donor (‘HMEJ’) predicted to favor homology-mediated end joining^18,19^ (Fig. 2A). Both knockin donors were provided on plasmids with the knockin sequence flanked by ~800bp homology arms, the length of which did not significantly affect results within a range of 500 to 1500 bp (data not shown). The HMEJ donor differed from the HR donor by the insertion of gRNA Binding Sites (GRBS) flanking the homology arms, which are cleaved by Cas9 to create linear dsDNA donors in cells. The orientation of the GRBSs did not have significant effects on editing performance (Supplementary Fig. S2).

**Fig. 2.**
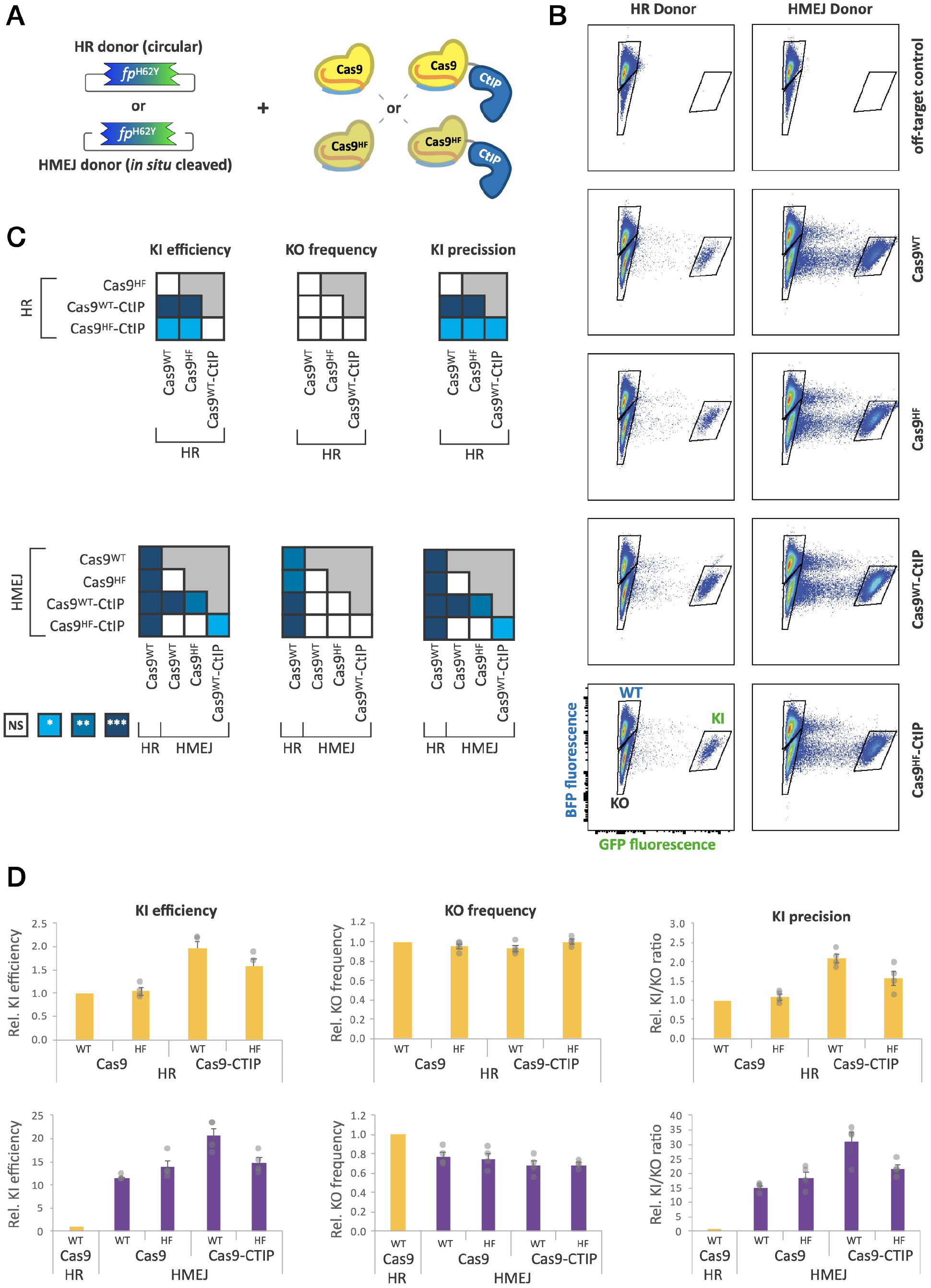
Cas9-CtIP fusion and HMEJ donors additively improve knockin precision. **(A)** Schema illustrating the combinatorial parameters of editing agents: dsDNA donor template HR or HMEJ variants in combination with Cas9 WT or HiFi variants, alone or fused to CtIP. **(B)** Flow cytometry plots of *HEK:BFP* cells 7 days after transient transfection with gRNA, Cas9, and donor plasmids. GFP fluorescence is shown on the x-axis (log), and BFP fluorescence on the y-axis (log). Quantification gates are indicated on the plots for BFP^+^ (WT), GFP^+^ (KI), and dark (KO) cells. **(D)** Quantification of flow data representing normalized knockin efficiency (%GFP+), knockout frequency (% dark), and knockin precision. Cas9 variants with HR (top row, yellow) or HMEJ (bottom row purple) donors were measured. Mean values from individual experiments (n ≥3) were normalized to those of the Cas9 WT with HR donor condition and presented as the mean ± SEM. **(C)** Heatmaps displaying statistical significance of pairwise comparisons of editing agent performance, calculated using one-way ANOVA with Tukey’s multiple comparison test and single pooled variance (* P<0.05; ** P<0.01; *** P < 0.001). Cas9-CtIP fusion with HMEJ donor outperforms Cas9 with HR donor in knockin precision by over 30-fold.

Transfection of HEK:BFP cells with Cas9 variant, donor template, and the *BFP* gRNA led to the emergence of both knockin (GFP^+^) and knockout (dark) populations for all combinations of donors and Cas9 variants, while transfection with off-target control gRNA resulted in no GFP^+^ cells (Fig. 2B). Among Cas9 variants tested, Cas9^WT^-CtIP performed with a roughly 2-fold improvement in knockin efficiency compared to Cas9^WT^, regardless of dsDNA donor variant, consistent with previous reports.^17,20,21^ Interestingly, CtIP fusion did not significantly improve efficiency and precision for single strand oligodeoxynucleotide DNA (ssODN) donors (Supplementary Fig. S3). Cas9^HF^ variants were generally not significantly different from Cas9^WT^, although Cas9^HF^-CtIP showed a 1.6-fold improvement in knockin efficiency specifically with the HR donor. Knockout rates did not significantly differ among the Cas9 variants (Fig. 2C), and thus, the KI/KO ratios (knockin precision) mirror the differences seen in knockin efficiency (Fig. 2D).

In contrast to the Cas9 variants, donor architecture impacted knockin efficiency as well as knockout rates. Across all four Cas9 variants, the HMEJ donor showed a 9- to 13-fold increase in knockin efficiency compared to the corresponding HR combination. Of the combinations of Cas9 and donors tested, Cas9^WT^-CtIP in conjunction with the HMEJ donor resulted in the highest rate of knockin, 24-fold higher than Cas9^WT^ with the HR donor. Regardless of Cas9 variant, the HMEJ donor showed a 22-30% reduction in gene disruption, which, together with the improved knockin efficiency, enhanced the precision by about 15-fold relative to the HR donor template permutations (Supplementary Fig. S2). Interestingly, these data suggest the independent and additive contributions of both CtIP fusion (~2-fold) and the *in situ* cleaved HMEJ donors (~11-fold) to knockin efficiency. Taken together, these results demonstrate that combinatorial optimization of both Cas9 and donor DNA can significantly shift the balance away from error-prone repair, with the combination of Cas9^WT^-CtIP (henceforth “Cas9-CtIP”) and HMEJ donors giving the highest efficiency and precision in HEK cells.

### Compound fusions on Cas9 improve editing performance

Several groups have independently demonstrated that modulation of DNA repair pathways is an effective way to improve knockin efficiency and precision. ^17,21–28^ To build on the results of the 3-factor screening highlighting WT Cas9, CtIP fusion, and HMEJ donor as the best performing combination, we iterated the BFP-to-GFP screening platform with constant HMEJ donor and evaluated the impact of five candidate DNA repair protein domains (dn53BP1,^24–26,28^ TIP60,^24,27^ RNF169,^26^ Rad52,^21–23,25–27^ eRad18^27^) on editing efficacy and precision when fused N-terminally to Cas9 or Cas9-CtIP (Fig. 3A).

**Fig. 3.**
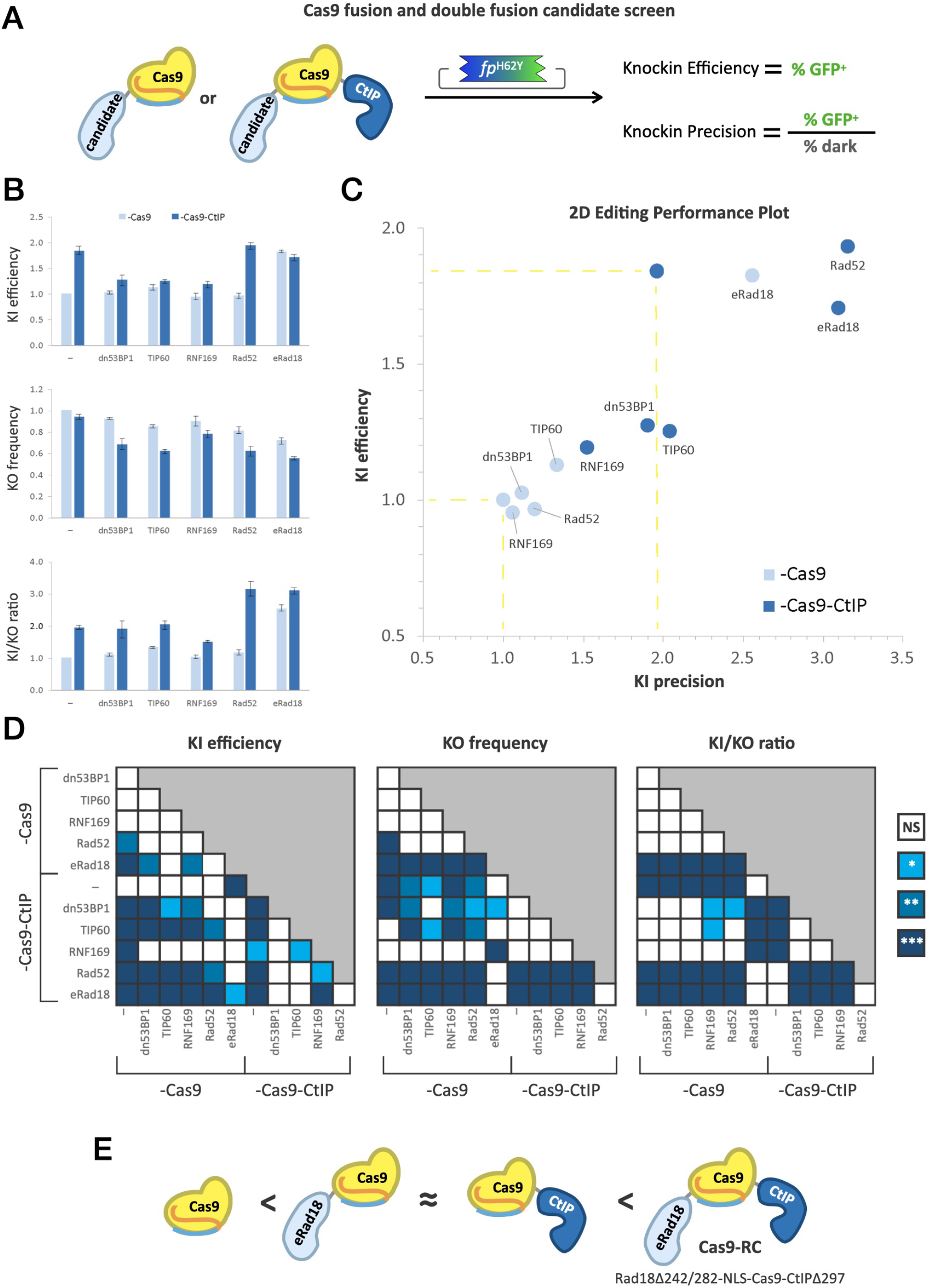
Iterative screening of novel Cas9 fusions and compound fusions with DNA-repair domains for increased editing performance. **(A)** Schema of the fusion screen. Candidate DNA-repair protein domains are fused N-terminally to either Cas9 alone or Cas9-CtIP and together with HMEJ donor assayed by BFP-to-GFP for knockin efficiency and precision. **(B)** Bar graphs showing quantification of relative knockin efficiency, knockout frequency, and knockin precision for Cas9 fusion (light blue) or Cas9-CtIP compound fusion (dark blue) with the listed DNA-repair protein domains. Values from individual experiments (n ≥ 3) were normalized to Cas9 only and presented as the mean ± SEM. **(C)** 2D editing performance plot comparing relative knockin efficiency and precision of Cas9 fusions (light blue) or Cas9-CtIP compound fusions (dark blue). Points with yellow dashed lines projecting to the axes correspond to Cas9 alone (normalization reference) and Cas9-CtIP. **(D)** Heatmaps showing statistical significance calculated using one-way ANOVA with Tukey’s multiple comparison test and single pooled variance (* P<0.05; ** P<0.01; *** P < 0.001). Data of replicate experiments are shown in Supplementary Fig. S5. **(E)** Schema illustrating comparative performance outcomes: novel fusion eRad18-Cas9 has equivalent performance to Cas9-CtIP. Novel compound fusion eRad18-Cas9-CtIP, henceforth named Cas9-RC, outperforms single fusions.

In the absence of CtIP, only eRad18 fusion to Cas9 significantly improved knockin efficacy, increasing it by 1.8-fold above Cas9 alone, similar to Cas9-CtIP. With the compound fusions, while addition of dn53BP1, TIP60, or RNF169 to Cas9-CtIP appeared to abrogate the effect of CtIP on knockin efficacy, fusion of Rad52 or eRad18 did not have a detrimental impact on efficiency (Fig. 3B-D).

Regarding unintended on-target knockout, dn53BP1-, TIP60-, and RNF169-fused Cas9 did not significantly differ from the Cas9-only control. Rad52 and eRad18 fusion, however, showed 18% and 28% reductions in knockout frequency, respectively (Fig. 3B-D). Interestingly, compound fusion of each of the five DNA repair proteins with CtIP led to significant reductions in the knockout rate, with eRad18, Rad52, and TIP60 showing the most pronounced decreases (45%, 38% and 38%, respectively). Despite these reductions in knockout rates, only Rad52 and eRad18 led to significant improvements in overall knockin precision. Without CtIP, eRad18 demonstrated a 2.5-fold increase in the knockin-to-knockout ratio, while combination of either Rad52 or eRad18 with CtIP led to a 3.1-fold increase in precision relative to Cas9 alone (Fig. 3B-D). These results show that specific combinations of DNA repair domains can function together to improve both the efficiency and precision of editing with Cas9.

### Novel compound fusion Cas9-RC increases knockin performance *in vivo*

The combinatorial screening and iterative optimizations of Cas9 agents yielded novel compound fusions of Cas9 with DNA repair protein domains that showed order-of-magnitude increases in knockin performance in cultured cells. Aiming to develop precision knockin agents for direct editing in vivo, we sought to assess the knockin efficacy of the best performing agents, Cas9^WT^ fused to eRad18 and CtIP (“Cas9-RC”) with an HMEJ donor, in an *in vivo* mouse model. In addition to C-terminal fusion of truncated human CtIP, Cas9-RC harbors an N-terminal fusion of an “enhanced” variant of human Rad18 protein, eRad18, which contains a deletion Δ242-282 of a putative DNA-binding domain shown to enhance homology-dependent repair (HDR) when co-expressed with Cas9.^27^ The total length of Cas9-RC is 2,172 amino acid residues (see Supplementary Table S1).

To test the knockin efficiency of Cas9-RC *in vivo*, we used *in utero* electroporation^10^ in the embryonic mouse brain^29^ (Fig. 4A) targeting integration of a 2A mCherry cassette at the 3’ end of the endogenous β-Actin (*ActB*) locus (Fig. 4B). A combination of four plasmids containing Cas9 or Cas9-RC, HMEJ donor with the 2A mCherry knockin, *ActB* gRNA, and a GFP transfection marker were electroporated into embryonic day 14.5 (E14.5) wild-type mice targeting progenitors of projection neurons of sensorimotor cortex. When assessed at postnatal day 7 (P7), electroporation of Cas9-RC led to an increase in mCherry^+^ knockin neurons compared to Cas9 (Fig. 4C).

**Fig. 4.**
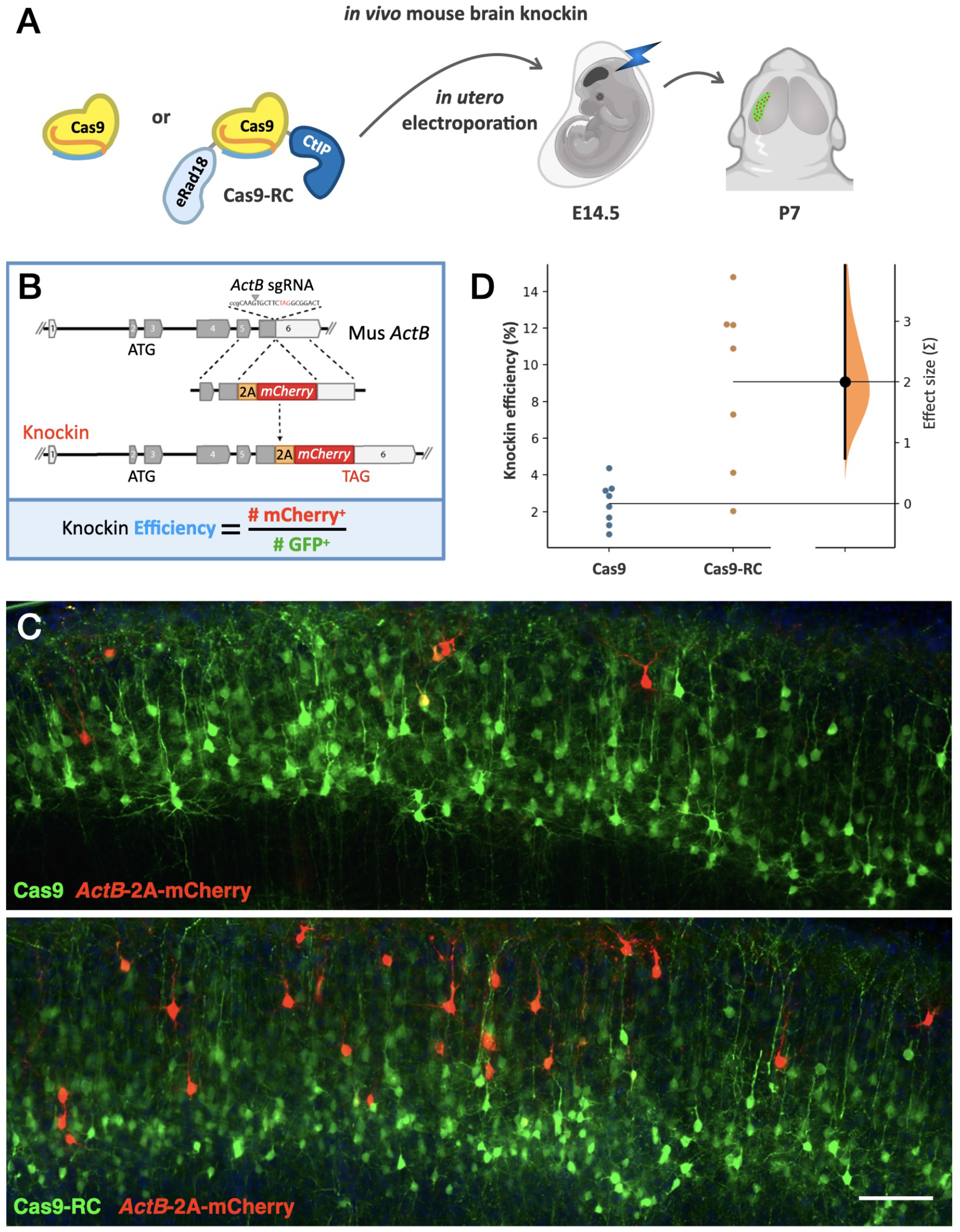
Cas9-RC enhances knockin efficiency *in vivo*. **(A)** Schema of the Cas9 and Cas9-RC agents compared for *in vivo* editing using *in utero* electroporation in the fetal mouse brain at embryonic day (E) 14.5, and analysis in the cerebral cortex at postnatal day (P) 7. **(B)** Gene editing donor template DNA targets the endogenous *ActB* locus to insert mCherry downstream of the β-actin coding sequence separated by a 2A. Knockin efficiency was quantified as the number of mCherry^+^ (knockin) neurons over the number of GFP^+^ (electroporated) neurons. **(C)** Representative fluorescence images of the cerebral cortex at P7 with electroporated neurons receiving Cas9 or Cas9-RC. Images show plasmid GFP (green) and genomic *ActB*-2A-mCherry (red) expression. Scale bar 100 μm. **(D)** Swarm plots showing quantification of *in vivo* knockin efficiency for Cas9 vs. Cas9-RC and effect size estimation. Points show means from each brain and are plotted on the left axis for both groups indicating knockin efficiency. The effect size on knockin efficiency of Cas9-RC versus Cas9 is plotted as a distribution on a floating axis on the right indicating standard deviations (Σ). The effect size estimated by unpaired Cohen’s d between Cas9 and Cas9-RC is 2.0 Σ (black dot), indicating a large effect size. The 95% confidence interval is 0.739 to 3.97 (vertical error bar). The P value is 0.0022.

*In vivo* knockin efficiency was calculated by comparing the number of mCherry^+^ knockin neurons to the number of GFP^+^ electroporated neurons (Fig. 4B). Electroporation of Cas9 resulted in an *in vivo* knockin efficiency of 2.4% (± 0.44%), while Cas9-RC yielded a 3.7-fold increase in performance averaging 9% (± 1.7%) *in vivo* knockin efficiency (Fig. 4C-D). These results demonstrate that Cas9-RC outperforms existing DSB repair-based editing agents with over three-fold increases in knockin performance both *in vitro* and *in vivo*.

## Discussion

In this study we sought to develop high-performance knockin tools by exploring combinations of DNA donor templates, variants of Cas9, and fusion of DNA repair protein domains. We identified novel Cas9 fusions and donor combinations that resulted in knockin with significantly improved metrics of editing efficiency and precision, *in vitro* and *in vivo*.

Our work establishes a standardized pipeline to optimize tools for precision genome editing. The BFP-to-GFP screening platform in HEK cells provides a high-throughput quantitative readout of the efficiency of correctly edited cells, while also reporting on the frequency of incorrectly edited cells. By simultaneously evaluating knockin and knockout rates, we identified combinations that optimized both efficiency (overall knockin rate) and precision (knockin rate vs. knockout rate). Knockin of a fluorescence cassette into the highly expressed β-Actin locus similarly enabled direct quantification of efficiency for large inserts at an endogenous gene locus *in vivo*.

We propose editing *efficiency* and *precision* as generalizable performance metrics for comparing genome editing agents. The ratiometric nature of these values make them versatile enough to be applied in a variety of biological systems regardless of whether the experimental outputs are sequences (i.e. Fig. 1C), surrogate cellular markers (i.e. Fig 2), or even function. These metrics incorporate all elements of the editing agent, including formulation (e.g. plasmid, RNP), editing modality (e.g. DSB repair, base editing, Prime editing), and delivery system (e.g. lipids, viral, nanoparticles), all of which may affect both outcomes. Efficiency and precision can be used as readouts when assessing individual components for holistic performance optimization, as we have done here. They can additionally be useful as common metrics to compare performance across distinct editing modalities, e.g., DSB repair vs Prime editing. By simultaneously assessing efficiency and precision across a variety of knockin tools, optimal agents can be selected based on the experimental need. For example, with *ex vivo* editing, one might prioritize efficiency if there are facile methods for *post hoc* selection of properly edited cells, whereas precision may be prioritized for contexts where edited cells cannot be selected, such as with *in vivo* editing.

Using this dual metric performance assessment, our study developed the high-performance DSB repair editor Cas9-RC (see Supplementary Discussion). When paired with HMEJ donor templates, Cas9-RC outperformed other knockin agents by over 30-fold in human cells and over 3-fold in the mouse brain. Importantly, Cas9-RC enables high performance for large genomic edits, such as fluorescent protein knockin. This complements parallel developments in base editors and Prime editing, which offer high performance but are limited to smaller edits. Ultimately, a diverse toolkit of precision editors will be useful to broaden the scope of *in vivo* editing applications. As presented in our study, having standardized platforms for quantitative comparison of new tools and novel combinations will further support efforts towards precision *in vivo* editing for both basic research and the development of future human therapeutics.

## Conclusion

Fusion of Cas9 to DNA repair protein domains can produce synergistic enhancements of knockin performance. Iterative high-throughput screening based on fluorescent protein conversion is an effective platform to assess knockin efficiency and precision for developing new editing agents. Cas9-RC is a new DSB repair genome editor demonstrating enhanced knockin performance *in vitro* and *in vivo*.

## Supporting information

Supplementary Discussion

## Acknowledgements

We are grateful for the technical and instrumentation support of the University of Maryland School of Medicine Center for Innovative Biomedical Resources through the Confocal Imaging Facility, the Flow Cytometry Facility, the Genomics Facility, and the Biostatistics Shared Service.

