## Supplementary Figures for "Cas9 fusions for precision *in vivo* editing"

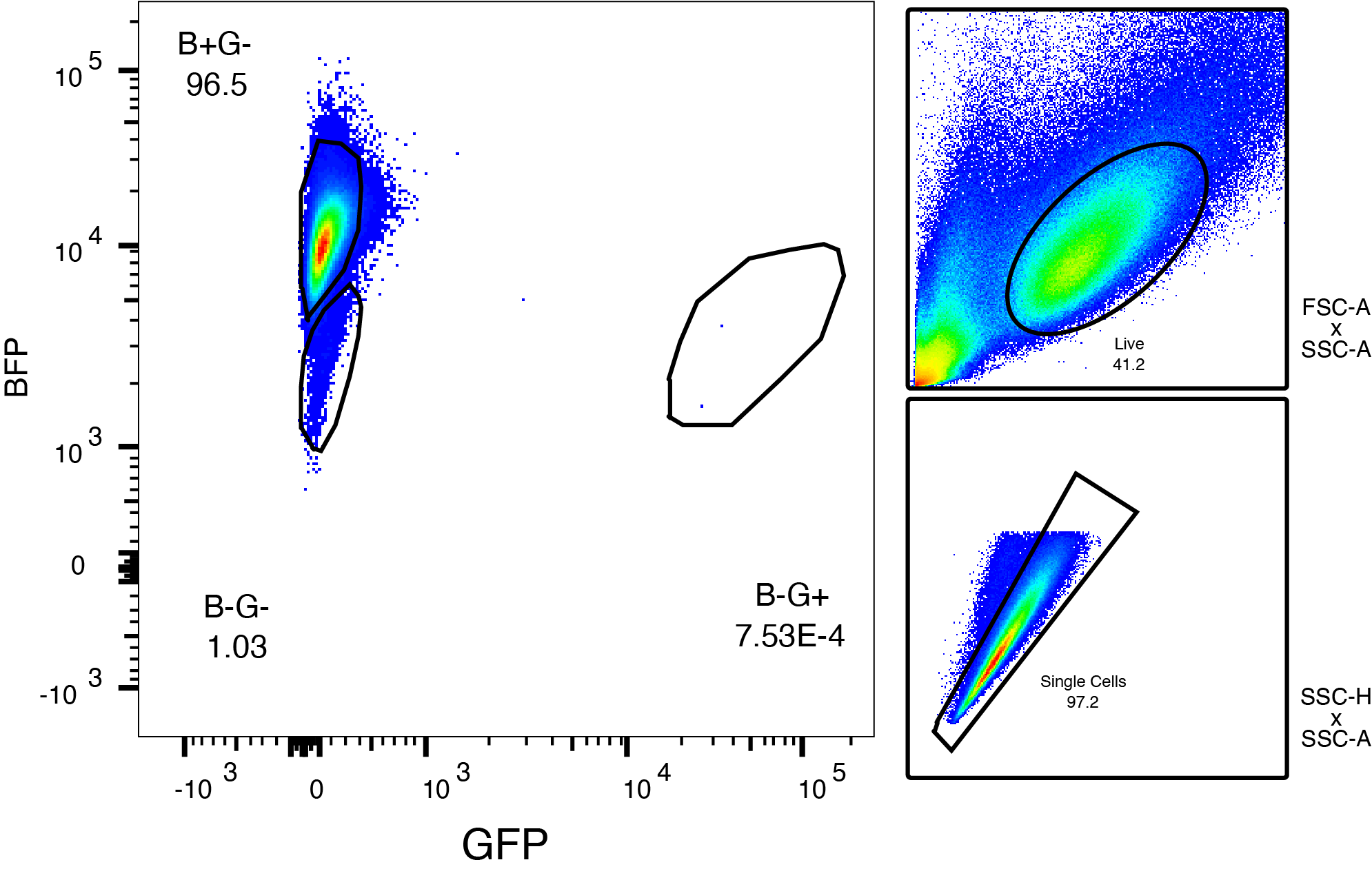

**Supplementary Fig. S1**. Gating strategy for flow cytometry analysis of editing in *HEK:BFP* cells. Live cells were first gated by size and granularity using FSC-A vs SSC-A and then singlets were gated using SSC-A vs SSC-H.

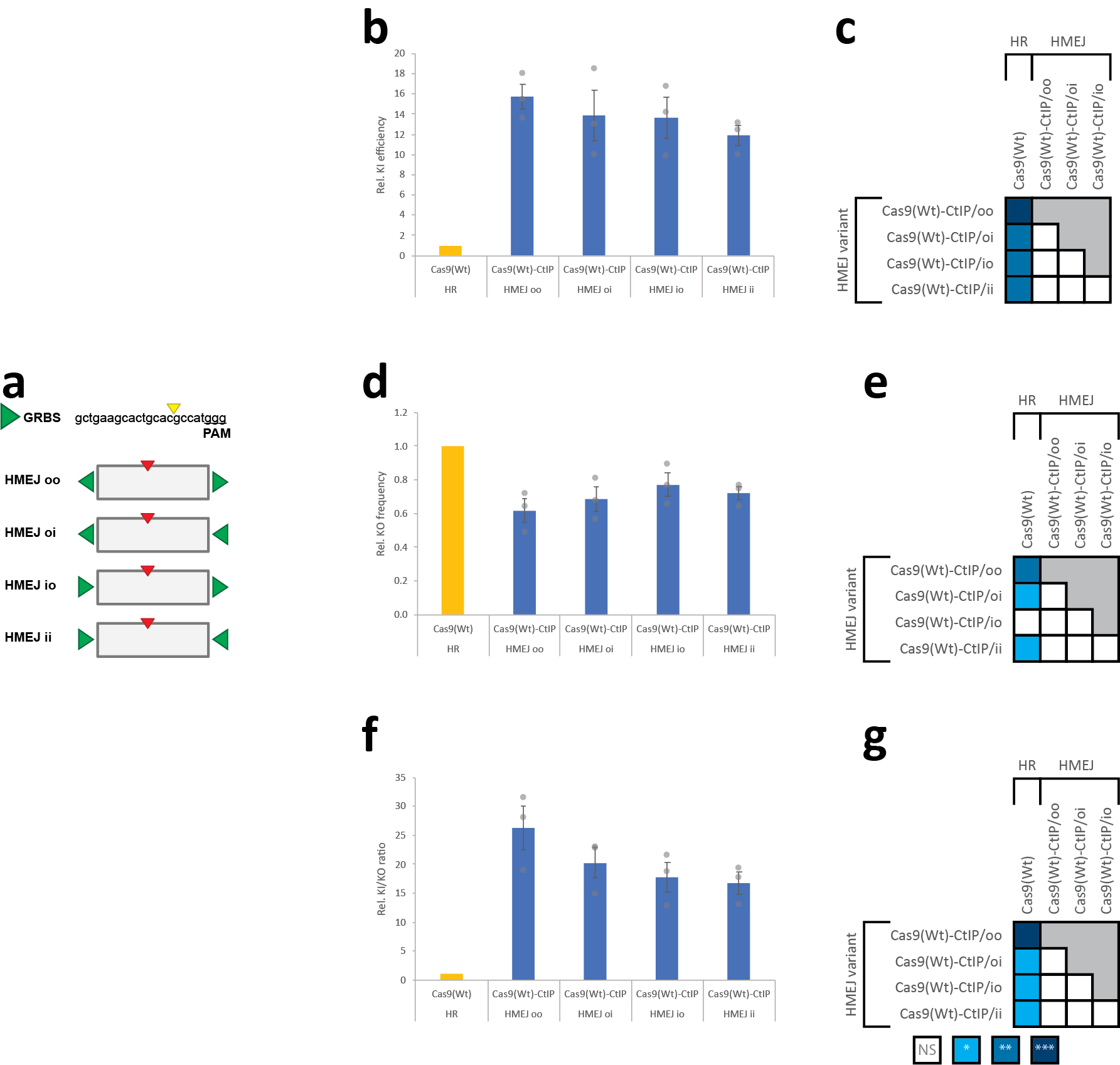

**Supplementary Fig. S2**. GRBS orientation does not impact efficiency or precision for HMEJ donors. **(a)** Schematic depicting asymmetry of BFP gRNA target sequence (GRBS) and HMEJ donors with all permutations of GRBS orientation. **(b,d,f)** Quantification of flow cytometry data from *HEK:BFP* cells 7 days after transient transfection indicating **(b)** KI efficiency (% GFP^+^) **(d)** KO efficiency (% dark) and **(f)** KI precission (KI/KO ratio) for HMEJ variants and Cas9(Wt)-CtIP. Values from individual experiments (n=3) were normalized to the Cas9(Wt)/HR donor condition and presented as the mean ± SEM. **(c,e,g)** Statistical significance was calculated using a one-way ANOVA with Tukey’s multiple comparison test, with a single pooled variance (*, P < 0.05; **, P < 0.01; ***, P < 0.001).

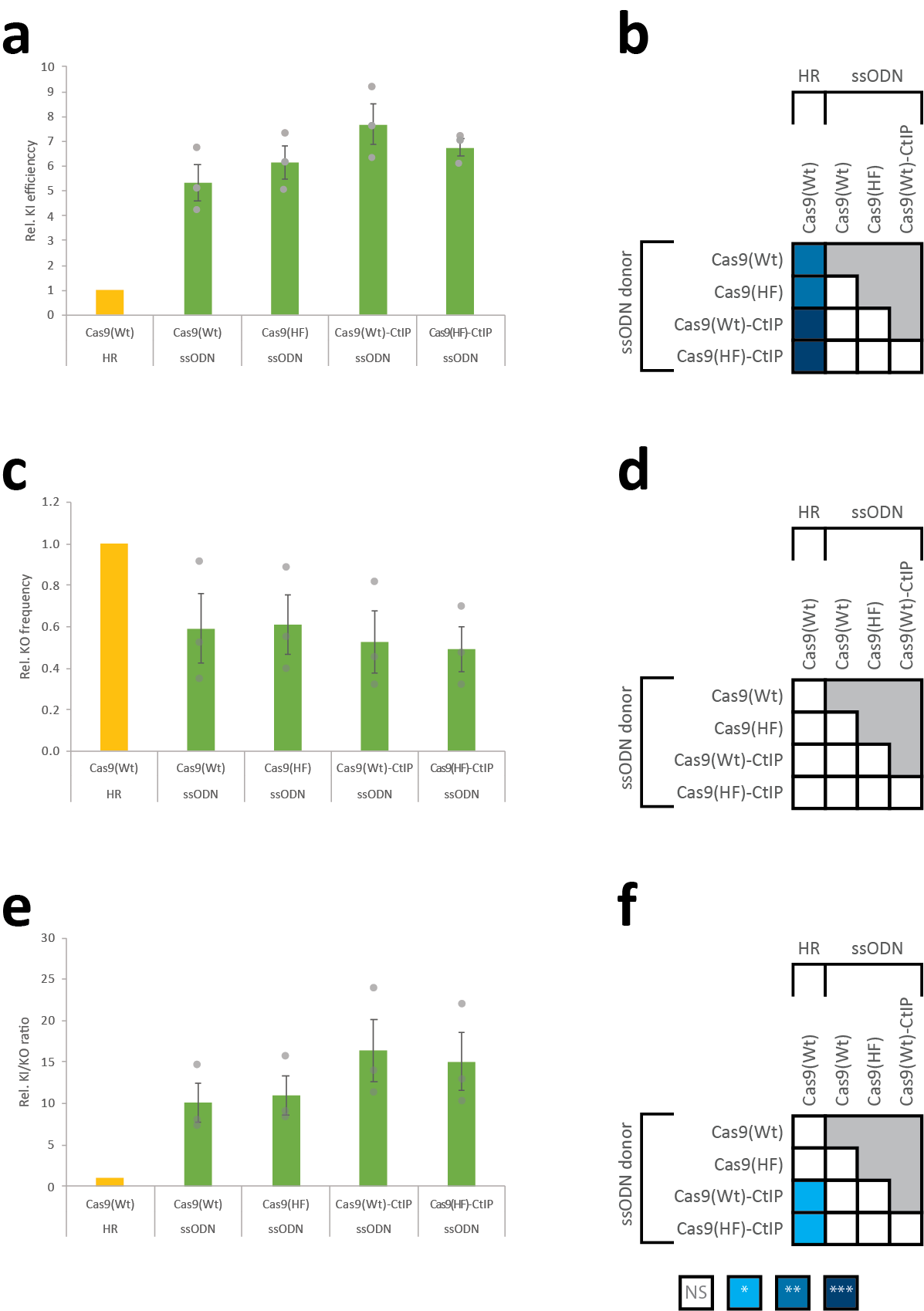

**Supplementary Fig. S3**. CtIP fusion does not significantly improve efficiency and precision for ssODN donors. **(a,c,e)** Quantification of flow cytometry data from *HEK:BFP* cells 7 days after transient transfection indicating **(b)** KI efficiency (% GFP^+^) **(d)** KO efficiency (%BFP^-^) and **(f)** KI/KO ratio for Cas9 variants with a ssODN donor. Values from individual experiments (n=3) were normalized to the Cas9(Wt)/HR donor condition and presented as the mean ± SEM. **(b,d,f)** Statistical significance was calculated using a one-way ANOVA with Tukey’s multiple comparison test, with a single pooled variance (*, P < 0.05; **, P < 0.01; ***, P < 0.001).

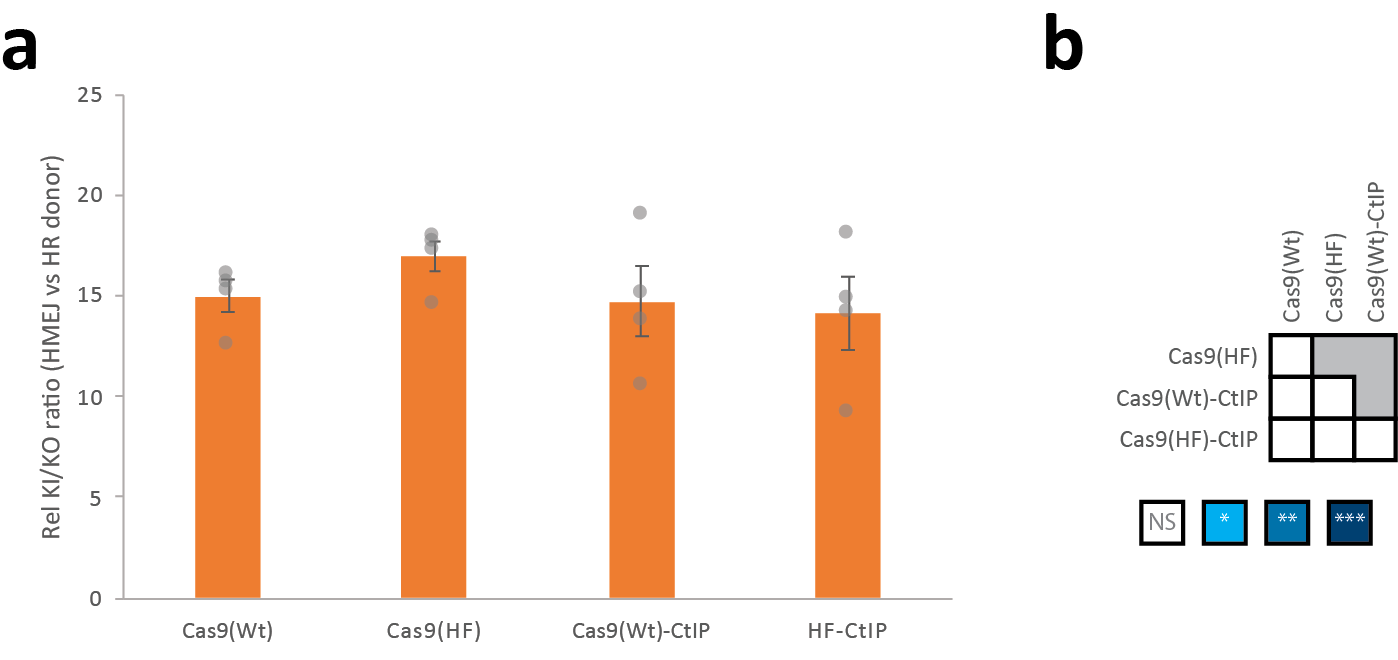

**Supplementary Fig. S4**. HMEJ donor independently improves precision across Cas9 variants. **(a)** Data related to Fig. 2 were analyzed to compare differences in precision between HR and HMEJ donors for the same Cas9 variant. **(b)** Statistical significance was calculated using a one-way ANOVA with Tukey’s multiple comparison test, with a single pooled variance. Differences between conditions were judged to be significant at P < 0.05 (*), P < 0.01 (**), and P < 0.001 (***). HMEJ outperforms HR donors by approximately 15-fold equivalently in all Cas9 iterations tested.

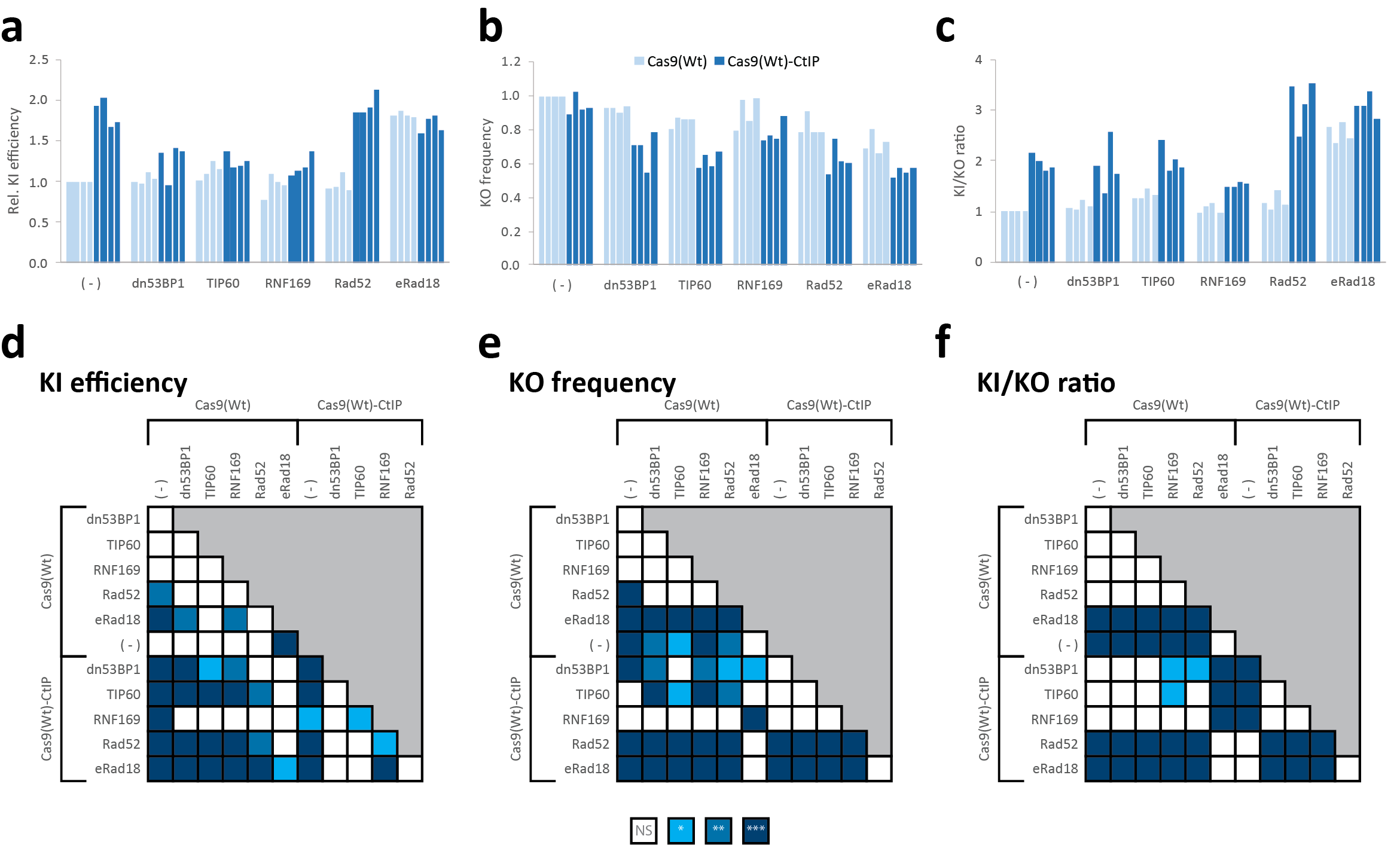

**Supplementary Fig. S5**. Biological replicate data and statistical analysis related to Fig. 3. **(a-c)** Quantification of flow cytometry data for biological replicates from HEK:BFP cells 7 days after transient transfection indicating **(a)** KI efficiency (% GFP^+^) **(b)** KO frequency (% dark) and **(c)** KI precision (KI/KO ratio) for Cas9 variants with HMEJ donor. **(d-f)** Statistical significance was calculated using a one-way ANOVA with Tukey’s multiple comparison test, with a single pooled variance. Differences between conditions were judged to be significant at P < 0.05 (*), P < 0.01 (**), and P < 0.001 (***).
