## Supplementary Discussion for "Cas9 fusions for precision *in vivo* editing"

We explored various formats of knockin donor DNA, including plasmid DNA as well as in situ linearized DNA, where the homologous donor sequence is cut out of a plasmid by Cas9 in the cell. While such donors can be challenging to construct, ssDNA donors can be synthesized commercially as single strand oligodeoxynucleotide (ssODN) donors. However, unlike double strand DNA (dsDNA) donors, which likely engage homology-dependent repair pathways, ssODN donors mediate editing through synthesis-dependent strand annealing (SDSA) and ssDNA incorporation. In our hands, ssODN donors did not perform as well as HMEJ DNA donors (Supplementary Fig. S3), which are predicted to engage the HMEJ repair pathway. Thus, depending on the cell context, different donors can offer improvements in knockin efficiency depending on the balance of DNA repair pathways in the target cell.

When paired with an *in situ* linearized dsDNA donor, fusion of CtIP to wild-type Cas9 led to a 20-fold increase in knockin efficiency and 30-fold increase in editing precision in vitro (Fig. 2). Interestingly, our data indicate independent and additive contributions from both the CtIP fusion (approximately 2-fold increase) and the linear HMEJ donor (over 10-fold increase) to knockin efficiency, as well as an independent effect of the HMEJ donor on reducing knockout by over 25%. These observations support the previously observed individual increases in efficiency,[^17^](https://sciwheel.com/work/citation?ids=4974064&pre=&suf=&sa=0&dbf=0)^,^[^19^](https://sciwheel.com/work/citation?ids=3630571&pre=&suf=&sa=0&dbf=0) and further demonstrate cumulative efficacy and reduction of knockout rates, leading to synergistic optimizations of knockin performance.

The independence of CtIP and HMEJ donor effects suggests that these two optimizations function through different, parallel mechanisms. Indeed, while the role of CtIP in multiple homology-mediated repair pathways is well characterized, it has been speculated that linear donors may also function through a number of different repair pathways.[^16^](https://sciwheel.com/work/citation?ids=978416&pre=&suf=&sa=0&dbf=0)^,^[^18^](https://sciwheel.com/work/citation?ids=185992&pre=&suf=&sa=0&dbf=0)^,^[^19^](https://sciwheel.com/work/citation?ids=3630571&pre=&suf=&sa=0&dbf=0)^,^[^21^](https://sciwheel.com/work/citation?ids=7226541&pre=&suf=&sa=0&dbf=0) Difference in DNA repair pathway utilization across cell types and developmental stages may underlie the quantitative variability in knockin efficiencies observed across host cell types assayed in this study, and underscores the challenge in translating *in vitro* results to *in vivo* application.

We validated the correspondence of the phenotypic fluorescence outputs from the BFP-to-GFP conversion assay with the genotype of edited cells using decomposition of Sanger sequence data. While this approach lacks the resolution of next-generation sequencing techniques, its low cost and low technical barrier make it an attractive method for many labs to evaluate the fidelity of gene editing approaches in their own genes and cells of interest. Importantly, these *in vitro* assays enable high-throughput characterization of knockin reagents, from which the optima are selected for evaluation *in vivo*. The *in utero* *electroporation* assay subsequently allows *in vivo* delivery of the same plasmids that were screened *in vitro*, circumventing packaging limits of viruses, which are problematic delivery vectors for large fusion proteins such as these. Combining high throughput *in vitro* transfection and high yield *in utero* electroporation into one workflow offers further continuity and standardization for assessment of editing performance, as well as iterative cycles of *in vitro* to *in vivo* optimization.

The observation of cumulative improvements on editing performance prompted us to further iterate the screening workflow to build upon the best performer of the previous cycle. We investigated fusion of additional DNA repair domains to Cas9-CtIP as a method to bias DSB repair towards knockin and away from deleterious repair. While small molecules and genetic manipulations can be used to skew the balance of repair pathways in cells, such global perturbations can result in cell toxicity that limit use *in vivo*. Directly fusing repair domains to Cas9 can obviate these toxicities by only modulating DNA repair locally at the target locus. Additionally, Cas9 fusions yield a single protein that is amenable to ribonucleoprotein delivery approaches.

We screened repair domains to fuse to Cas9 based on previous reports highlighting their impact on DNA repair outcomes. [^17^](https://sciwheel.com/work/citation?ids=4974064&pre=&suf=&sa=0&dbf=0)^,^[^21^](https://sciwheel.com/work/citation?ids=7226541&pre=&suf=&sa=0&dbf=0)^,^[^23^](https://sciwheel.com/work/citation?ids=6457074&pre=&suf=&sa=0&dbf=0)^-[28](https://sciwheel.com/work/citation?ids=7140365&pre=&suf=&sa=0&dbf=0)^ We found that N-terminal fusion of Rad52 and eRad18 improved the knockin precision of Cas9. Rad52 is a ssDNA-binding protein that mediates DNA repair both through the single-strand annealing (SSA) pathway in a Rad51-independent manner, as well as through homologous recombination in a Rad51-dependent manner. Despite this however, in our study, Rad52 did not significantly impact the efficiency of knockin whether in the presence or absence of CtIP. Instead, Rad52 fused to Cas9 alone reduced the frequency of knockout by 18%, and when combined with CtIP fusion led to an even greater 34% reduction. The *Saccharomyces cerevisiae* ortholog (ScRad52) may be more a potent knockin enhancer than the human variant that we used in this study, and it would be interesting to similarly characterize this related protein.[^22^](https://sciwheel.com/work/citation?ids=3102958&pre=&suf=&sa=0&dbf=0)^,[23](https://sciwheel.com/work/citation?ids=6457074&pre=&suf=&sa=0&dbf=0)^

Unlike Rad52, fusion of eRad18 to Cas9 showed comparable increases in knockin rates with fusion of Cas9 to CtIP. Consistent with this effect, eRad18 was previously demonstrated to stimulate homology-dependent repair (HDR) by inhibiting 53BP1 localization to DSBs. eRad18 further resulted in 24% less knockout when combined with CtIP in double fusions where Cas9 is flanked by the two optimized DNA repair domains (Cas9-RC). In all, our approach further supports the utility of biologically inspired, synthetic fusions for expanding the genome editing toolkit.
